# How macroecology affects macroevolution: the interplay between extinction intensity and trait-dependent extinction in brachiopods

**DOI:** 10.1101/523811

**Authors:** Peter D. Smits

**Affiliations:** University of California – Berkeley, Berkeley, California 94720

## Abstract

Selection is the force behind differences in fitness, with extinction being the most extreme example of selection. Modern experiments and observations have shown that average fitness and selection strength can vary over time and space. This begs the question: as average fitness increases, does selection strength increase or decrease? The fossil record illustrates how extinction rates have varied through time, with periods of both rapid and slow species turnover. Using Paleozoic brachiopods as a study system, I developed a model to understand how the average taxon duration (i.e. fitness) varies over time, to estimate trait-based differences in taxon durations (i.e. selection), and to measure the amount of correlation between taxon fitness and selection. I find evidence for when extinction intensity increases, selection strength on geographic range also increases. I also find strong evidence for a non-linear relationship between environmental preference for epicontinental versus open-ocean environments and expected taxon duration, where taxa with intermediate preferences are expected to have greater durations than environmental specialists. Finally, I find that taxa which appear more frequently in epicontinental environments will have a greater expected duration than those taxa which prefer open-ocean environments. My analysis supports the conclusions that as extinction intensity increases and average fitness decreases, as happens during a mass extinction, the trait-associated differences in fitness would increase. In contrast, during periods of low extinction intensity when fitness is greater than average, my model predicts that selection associated with geographic range and environmental preference would decrease and be less than average.

## Introduction

Selection is the force behind differences in fitness, with the most extreme example of selection being extinction. Modern experiments and paleontological analyses have demonstrated that selection strength and fitness can vary over time and space. An unanswered question in macroevolution is if and how species fitness and selection covary; does the strength of selection change as average fitness also changes? The fossil record demonstrates that extinction risk has varied continuously over time, from periods of low average extinction rate to very high extinction rates (e.g. mass extinctions) (Foote, 2000*a*,*b*, 2001). Paleontological analyses have also demonstrated trait-based differences in extinction risk among taxa (Jablonski, 2008). Conceptually, extinction is the ultimate manifestation of selection, as we would expect a taxon with a beneficial trait to persist longer than a similar taxon without that trait due to selection (Jablonski, 2008; Rabosky and McCune, 2010; Raup, 1994; Stanley, 1975). Thus, the expected duration of a species can be conceived of as a measure of a species’ fitness (Cooper, 1984); meaning that trait-associated differences in species fitness are species selection (Rabosky and McCune, 2010).

In order to test for an association between extinction intensity and extinction selectivity, extinction rate and trait-based differences in extinction rate need to be estimated. Previous work has approached this problem by estimating the extinction intensity and selectivity at different points in time, or for different origination cohorts independently and then comparing those estimates (Payne et al., 2016). I find this approach problematic for a few reasons. Modeling each time point or cohort independently does not use all of the information present in the data, and those estimates are only based on the data from that time point. A hierarchical/mixed-effect modelling approach leverages all data across time points or cohorts by partially pooling information across each of the time-points or cohorts. The resulting parameter estimates have better behaved posteriors (e.g. smaller credible intervals) and limit overly optimistic parameter estimates by weighing those estimates relative to the amount of data associated with each time point or cohort (Gelman et al., 2013). The partial pooling in hierarchical/mixed-effect models also mitigates the effects of complete separation, which normally prevents parameter estimates for some time points or cohorts (Gelman et al., 2013; Payne et al., 2016). Finally, treating each time point or cohort as statistically independent means that any and all post-hoc analyses are at risk of false positive results due to multiple comparisons issues (Gelman et al., 2013; Gelman and Hill, 2007); hierarchical/mixed-effect models ameliorate this problem as possible comparisons are modeled simultaneously.

Jablonski (1986) observed that for bivalves at the end-Cretaceous mass extinction event, previous trait-associated differences in survival no longer mattered except in the case of geographic range. Based on this evidence, Jablonski (1986) proposed the idea of “macroevolutionary regimes:” that mass extinction and background extinction are fundamentally different processes. However, based on estimates of extinction rates over time, there is no evidence of there being two or more “types” of extinction (Wang, 2003). Instead, extinction rates for marine invertebrates are unimodal with continuous variation. This disconnect between the qualitative differences of macroevolutionary modes and the observation of continuous variation in extinction rates implies the possibility of a relationship between the strength of selection (extinction intensity) and the association of traits and differences in fitness (extinction selectivity) (Payne et al., 2016). As extinction intensity increases, what happens to extinction selectivity? How do trait-associated differences in fitness change as average extinction rate changes over time?

Here I develop a statistical model describing the relationship between brachiopod taxon durations and multiple functional taxon traits in order to understand the relationship between extinction intensity and selectivity over time. Trait-dependent differences in extinction risk should be associated with differences in taxon duration (Cooper, 1984; Rabosky and McCune, 2010). Brachiopods are an ideal group for this study as they have an exceptionally complete fossil record (Foote, 2000*b*; Foote and Raup, 1996). I focus on the brachiopod record from the post-Cambrian Paleozoic, from the start of the Ordovician, approximately 485 million years ago (Mya), through the end Permian (approximately 252 Mya) as this represents the time of greatest global brachiopod diversity and abundance (Alroy, 2010).

The analysis of taxon durations, the time from a taxon’s origination to its extinction, falls under the purview of survival analysis, a field of applied statistics commonly used in health-care and engineering (Klein and Moeschberger, 2003), that has a long history in paleontology (Crampton et al., 2016; Simpson, 1944, 1953; Smits, 2015; Van Valen, 1973, 1979). I adopt a hierarchical Bayesian modeling approach (Gelman et al., 2013; Gelman and Hill, 2007) in order to unify the previously distinct dynamic and cohort paleontological survival approaches (Baumiller, 1993; Crampton et al., 2016; Ezard et al., 2012; Foote, 1988; Raup, 1975, 1978; Simpson, 2006; Van Valen, 1973, 1979).

While estimation of trait-dependent speciation and extinction rates from phylogenies of extant taxa has become routine (Fitzjohn, 2010; Goldberg et al., 2011, 2005; Maddison et al., 2007; Rabosky et al., 2013; Stadler, 2011, 2013; Stadler and Bokma, 2013), there are two major ways to estimate trait-dependent extinction: analysis of phylogenies and analysis of the fossil record. These two directions, phylogenetic comparative and paleobiological, are complementary and intertwined in the field of macroevolution (Hunt and Rabosky, 2014; Jablonski, 2008; Rabosky and McCune, 2010). In the case of extinction, analysis of the fossil record has the distinct advantage over phylogenies of only extant taxa because extinction is observable; this means that extinction rates can be directly estimated (Liow et al., 2010; Quental and Marshall, 2009; Rabosky, 2010). The approach used here is thus complementary to the analysis of trait-dependent extinction based phylogenetic structure.

### Factors affecting brachiopod survival

Conceptually, taxon survival can be considered an aspect of “taxon fitness” (Cooper, 1984; Palmer and Feldman, 2012). Traits associated with taxon survival are thus examples of species (or higher-level) selection, as differences in survival are analogous to differences in fitness. The traits analyzed here are all examples of emergent and aggregate traits (Jablonski, 2008; Rabosky and McCune, 2010). Emergent traits are those which are not measurable at a lower level (e.g. species are a higher level aggregate of individual organism) such as global geographic range, or fossil sampling rate. Aggregate traits, like body size or environmental preference, represent the average of a shared trait across all members of a lower level.

Geographic range is widely considered the most important biological trait for estimating differences in extinction risk at nearly all times, with large geographic range conferring low extinction risk (Finnegan et al., 2012; Harnik et al., 2012; Jablonski, 1986, 1987, 2008; Jablonski and Roy, 2003; Payne and Finnegan, 2007). This relationship is expected even if extinction is a completely random process. Because of its importance and size, geographic range was analyzed as a covariate of extinction risk with the initial assumption that a taxon with greater than average geographic range would have a lower than average extinction risk. The effect size of geographic range on extinction acts as a baseline for comparing the strength of selection associated with the other covariates.

Epicontinental seas are a shallow-marine environment where the ocean has spread over the continental interior or craton with a depth typically less than 100 meters. In contrast, open-ocean coastline environments have much greater variance in depth, do not cover the continental craton, and can persist during periods of low sea level (Miller and Foote, 2009). This hypothesis is that taxa which favor epicontinental seas would be at a greater extinction risk during periods of low sea levels, such as during glacial periods, than environmental generalists or open-ocean specialists. Epicontinental seas were widely spread globally during the Paleozoic (approximately 541-252 Mya) but declined over the Mesozoic (approximately 252–66 My) and have nearly disappeared during the Cenozoic (approximately 66–0 My) as open-ocean coastlines became the dominant shallow-marine setting (Johnson, 1974; Miller and Foote, 2009; Peters, 2008; Sheehan, 2001). Taxa in epicontinental environments could also have a greater extinction susceptibility than taxa in open-ocean environments during anoxic events or other major changes to water chemistry due to limited water circulation from the open-ocean into epicontiental seas (Peters, 2007). Similarly, if there is a major and sudden change to water chemistry in a single epicontinental sea, the sluggish water flow into and out of that sea would most likely not affect other epicontinental seas leading to local extirpation but not global extinction.

Miller and Foote (2009) demonstrated that, during several mass extinctions, taxa associated with open-ocean environments tended to have a greater extinction risk than those taxa associated with epicontinental seas. During periods of background extinction, however, they found no consistent difference in extinction risk between taxa favoring either environment. Miller and Foote (2009) hypothesize that open-ocean taxa may have a greater extinction rate because these environments would be more strongly affected by poisoning of the environment from impact fallout or volcanic events because water circulates at a higher rate and in greater volume in open-ocean environments compared to the relatively more sluggish ciruclation in epicontinental environments. These two environment types represent the primary identifiable environmental dichotomy observed in ancient marine systems (Miller and Foote, 2009; Sheehan, 2001). Given these findings, I would hypothesize that as average extinction risk increases, the difference in risk associated with open-ocean environments versus epicontinental environments should generally increase.

Because environmental preference is defined here as the continuum between occurring exclusively in open-ocean environments versus epicontinental environments, intermediate values are considered “generalists” in the sense that they favor neither end-member. A long-standing hypothesis is that generalists or unspecialized taxa will have greater survival than specialists (Baumiller, 1993; Liow, 2004, 2007; Nürnberg and Aberhan, 2013, 2015; Simpson, 1944; Smits, 2015). Because of this, the effect of environmental preference was modeled as a quadratic function, where a concave-down relationship between preference and expected duration indicates that generalists are favored over specialists end-members. Importantly, this approach does not “force” a non-linear relationship and only allows one if the second-order term is non-zero.

Body size, measured as shell length, is also considered as a trait that may potentially influence extinction risk (Harnik, 2011; Payne et al., 2014). Body size is a proxy for metabolic activity and other correlated life history traits (Payne et al., 2014). Harnik et al. (2014) analyzed the effect of body size selectivity in Devonian brachiopods in both phylogenetic and non-phylogenetic contexts and found that that body size was not associated with differences in taxonomic duration. However, there are some bivalve subclades for which body size can be as important a factor as geographic range size in determining extinction risk (Harnik, 2011). Given these results, I expect that if body size has any effect on brachiopod taxonomic survival, it will be very small.

It is well known that, given the incompleteness of the fossil record, the observed duration of a taxon is an underestimate of that taxon’s true duration (Alroy, 2014; Foote and Raup, 1996; Liow and Nichols, 2010; Solow and Smith, 1997; Wagner and Marcot, 2013; Wang and Marshall, 2004). Because of this, the concern is that a taxon’s observed duration may reflect its relative chance of being sampled and not any of the effects of the covariates of interest. In this case, for sampling to be a confounding factor there must be consistent relationship between the quality of sampling of a taxon and its apparent duration (e.g. greater sampling, longer duration). If there is no relationship between sampling and duration then interpretation can be made clearly; while observed durations are obviously truncated true durations, a lack of a relationship would indicate that the amount and form of this truncation is not a major determinant of the taxon’s apparent duration. By including sampling as a covariate in the model, this effect can be quantified and can be taken into account when interpreting the estimates of the effects of the other covariates.

## Methods

The brachiopod dataset analyzed here was sourced from the Paleobiology Database (http://www.paleodb.org) which was limited to Brachiopods as defined by the higher taxonomic groups Rhychonelliformea: Rhynchonellata, Chileata, Obolellida, Kutorginida, Strophomenida, and Spiriferida. Additionally, samples were limited to those which originated after the Cambrian but before the Triassic. Temporal, stratigraphic, and other relevant occurrence information used in this analysis was also downloaded from the same source. Analyzed occurrences were restricted to those with paleolatitude and paleolongitude coordinates, being assigned to either epicontinental or open-ocean environment, and belonging to a genus present in the body size dataset (Payne et al., 2014). Epicontinental versus open-ocean assignments for each fossil occurrence are based on those from previous analyses (Foote and Miller, 2013; Miller and Foote, 2009; Ritterbush and Foote, 2017). These filtering criteria are very similar to those from Foote and Miller (2013) with the additional constraint to only those taxa which occur in the body size data set from Payne et al. (2014). In total, 1130 genera were included in this analysis.

Fossil occurrences were analyzed at the genus level, a common practice for paleobiological, macroevolutionary, and macroecological studies, and this is especially the case for marine invertebrates (Alroy, 2010; Eronen et al., 2011; Foote and Miller, 2013; Harnik et al., 2012; Kiessling and Aberhan, 2007; Miller and Foote, 2009; Nürnberg and Aberhan, 2013, 2015; Payne and Finnegan, 2007; Ritterbush and Foote, 2017; Simpson and Harnik, 2009; Vilhena et al., 2013). While species diversity dynamics are frequently of much greater interest than those of higher taxa (though see Foote 2014; Hoehn et al. 2015), the nature of the fossil record makes accurate, precise, and consistent taxonomic assignments at the species level difficult for all occurrences. To ensure a minimum level of confidence and accuracy in the data, I analyzed genera as opposed to species. Additionally, when species and genera can be compared, they often yield similar results (Foote et al., 2007; Jernvall and Fortelius, 2002; Roy D. & Valentine, I. W., 1996). Importantly, it is also possible that genera represent coherent biological units as there is evidence for congruence between morphologically and genetically defined genera of molluscs and mammals (Jablonski and Finarelli, 2009).

Genus duration was calculated as the number of geologic stages from first appearance to last appearance, inclusive. Durations were based on geologic stages as opposed to millions of years because of the inherently discrete nature of the fossil record. Dates are not assigned to individual fossils themselves; rather fossils are assigned to a geological interval which represents some temporal range. In this analysis, stages are effectively irreducible temporal intervals in which taxa may occur. Genera with there last occurrence in or after Changhsingian stage (e.g. the final stage of the study interval) were right-censored. Censoring in this context indicates that the genus was observed up to a certain age, but that its ultimate time of extinction is unknown (Klein and Moeschberger, 2003).

The covariates of duration included in this analysis are geographic range size (*r*), environmental preference (*v*, *v*^2^), shell length (*m*), and sampling (*s*).

The geographic range of a genus was calculated as the number of occupied grid cells from a gridded map of all contemporaneous occurrences. First, the paleolatitude-paleolongitude coordinates for all occurrences were projected onto an equal-area cylindrical map projection. Each occurrence was then assigned to one of the cells from a 70 × 34 regular raster grid placed on the map. Each grid cell represents approximately 250,000 km^2^. The map projection and regular lattice were made using shape files from http://www.naturalearthdata.com/ and the raster package for R (Hijmans, 2015). For each time interval a taxon’s geographic range is calculated from the ratio of cells occupied by that taxon divided by the total number of cells with any occurrences. For each taxon in each temporal bin, the relative occurrence probability of the observed taxa was calculated using the JADE method developed by Chao et al. (2015) which leverages the distribution of taxon occurrences to estimate their “true” geographic range. This method accounts for the fact that taxa with an occupancy of 0 cannot be observed, which means that occupancy follows a truncated Binomial distribution. This correction is critical when comparing occupancies from different times with different geographic sampling. After occurence probability is calculated for all taxa for each temporal bin in which they occur, I calculated mean occurrence probability of each taxa. This final value is my proxy for the geographic range of a taxon.

Environmental preference is a descriptor of whether and by how much a taxon prefers epicontinental to open-ocean environments. This approach presents environmental preference as a continuum from exclusive occurrence at the ends and equal occurrences in the middle. My measure of environmental preference is derived from the number of epicontinental or open-ocean observations of a taxon compared to the total number of epicontinenal or open-ocean observations that also occurred during time intervals shared with that taxon. Mathematically, environmental preference was defined as probability of observing the ratio of epicontinental occurrences to total occurrences (θ_*i*_ = *e_i_*/*E_i_*) or greater given the background occurrence probability 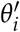 as estimated from all other taxa occurring at the same time 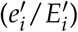. This measure of environmental preference is expressed

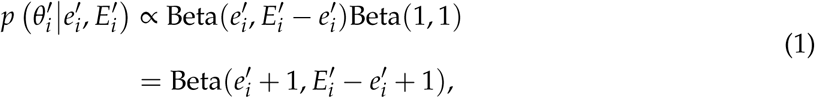

where *v* is the percent of the distribution defined in equation 1 as less than or equal to θ_*i*_. The Beta distribution is used here because it is a continuous distribution bounded at 0 and 1, which is idea for modeling percentages.

Body size, measured as shell length, was sourced directly from Payne et al. (2014). These measurements were made from brachiopod taxa figured in the *Treatise on Invertebrate Paleontology* (Williams et al., 2007).

The sampling probability for individual taxa, called *s*, was calculated using the standard gap statistic (Foote, 2000*a*; Foote and Raup, 1996). The gap statistic is calculated as the number of stages in which the taxon was sampled exempting its first and last stages. Because taxa that were right-censored only have a first appearance, one was subtracted from the numerator and denominator instead of two. The inclusion of genus-specific sampling probability as a covariate are an attempt to mitigate the effects of the incompleteness of the fossil record on our ability to observe genus duration. The implications of this choice are discussed further later in the Discussion.

The minimum duration for which a gap statistic can be calculated is three stages, so I chose the impute the gap statistic for all observations with a duration of less than 3. Imputation is the “filling in” of missing observations based on the observed values (Gelman and Hill, 2007; Rubin, 1996).

Prior to analysis, geographic range was logit transformed and the number of samples was natural-log transformed; these transformations make these variables defined for the entire real line. Sampling probability was transformed as (*s*(*n −* 1) + 0.5)/*n* where *n* is the sample size as recommended by Smithson and Verkuilen (2006); this transformation shrinks the range of the data so that there are no values equal to 0 or 1. All covariates except for sampling were standardized by subtracting the mean from all values and dividing by twice its standard deviation, which follows Gelman and Hill (2007). This standardization means that the associated regression coefficients are interpretable as the expected change per 1-unit change in the rescaled covariates. Finally, *D* is defined as the total number of covariates, excluding sampling, plus one for the intercept term.

### Details of model

Hierarchical modelling is a statistical approach which explicitly takes into account the structure of the observed data (Gelman et al., 2013; Gelman and Hill, 2007). The units of study (e.g. genera) each belong to a single group (e.g. origination cohort). Each group is considered a realization of a shared probability distribution (e.g. prior) of all cohorts, observed and unobserved. The group-level parameters, or the hyperparameters of this shared prior, are themselves given (hyper)prior distributions and are also estimated like the other parameters of interest (e.g. covariate effects) (Gelman et al., 2013). The subsequent estimates are partially pooled together, where parameters from groups with large samples or effects remain large while those of groups with small samples or effects are pulled towards the overall group mean. All covariate effects (regression coefficients), as well as the intercept term (baseline extinction risk), were allowed to vary by group (origination cohort). The covariance between covariate effects was also modeled.

Genus durations were assumed to follow a Weibull distribution, which allows for age-dependent extinction (Klein and Moeschberger, 2003): *y ∼* Weibull(α, σ). The Weibull distribution has two parameters: scale σ and shape α. When α = 1, σ is equal to the expected duration of any taxon. α is a measure of the effect of age on extinction risk, where values greater than 1 indicate that extinction risk increases with age, and values less than 1 indicate that extinction risk decreases with age. Note that the Weibull distribution is equivalent to the exponential distribution when α = 1.

Data censoring and truncation are conditions where the value of interest (taxon duration) is only partially observed. There are a number of processes which can lead to either of these conditions: limited resolution, which leads to left-censoring or truncation; end of study interval, which leads to right censoring; and incomplete sampling, which can left-censor (short-lived taxa are less likely to be preserved at all) or right-censor (durations are truncated). In the case of the right-and left-censored observations mentioned above, the probability of those observations has a different calculation (Klein and Moeschberger, 2003). For right-censored observations, the likelihood is calculated *p*(*y|θ*) = 1 − *F*(*y*) = *S*(*y*), where *F*(*y*) is the cumulative distribution function. Taxa that existed for only a single stage were left-censored, which implies that that taxon went extinct at any point between 0 and 1 stages. In contrast to right-censored data, the likelihood of a left-censored observation is calculated from *p*(*y|θ*) = *F*(*y*). This censoring strategy improves model fit, as measured by WAIC and LOOIC, than treating these taxa as being fully observed.

The scale parameter σ was modeled as a regression following Kleinbaum and Klein (2005) with varying intercept, varying slopes, and the effect of sampling. This model is expressed

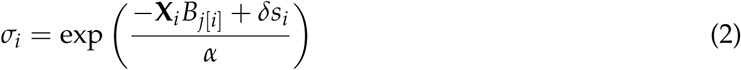

where *i* indexes across all observations from *i* = 1,…, *n*, *n* is the total number of observations, *j*[*i*] is the cohort membership of the *i*th observation for *j* = 1,…, *J*, *J* is the total number of cohorts, *X* is a *N × D* matrix of covariates along with a column of ones for the intercept term, *B* is a *J × D* matrix of cohort-specific regression coefficients, and δ is the regression coefficient for the effect of sampling *s*. δ does not vary by cohort.

Each of the rows of matrix *B* are modeled as realizations from a multivariate normal distribution with length *D* location vector µ and *J × J* covariance matrix Σ: *B_j_ ∼* MVN(µ, Σ). The covariance matrix was then decomposed into a length *J* vector of scales τ and a *J × J* correlation matrix Ω, defined Σ = diag(τ)Ωdiag(τ) where “diag” indicates a diagonal matrix.

The elements of µ were given independent normally distributed priors. The effects of geographic range size and the breadth of environmental preference were given informative priors reflecting the previous findings while the other parameters were given weakly informative priors which favor those covariates having no effect on duration. The correlation matrix Ω was given an LKJ distributed prior (Lewandowski et al., 2009) that slightly favors an identity matrix as recommended by Team (2017). These priors are defined

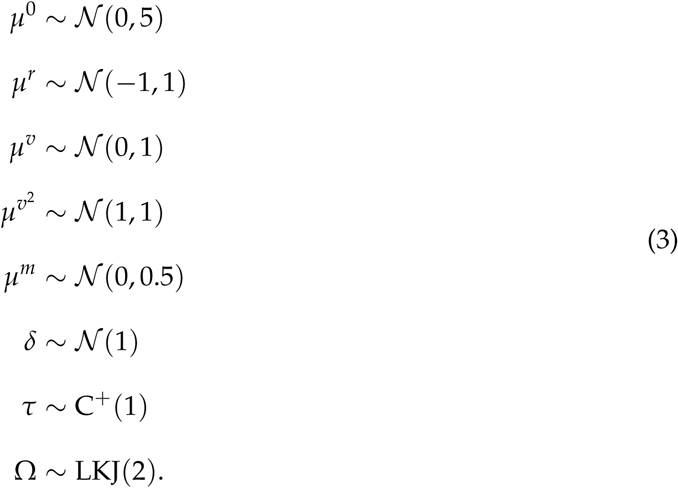

The log of the shape parameter α was given a weakly informative prior log(α) ∼ *N* (0, 1) centered at α = 1, which corresponds to the Law of Constant Extinction (Van Valen, 1973).

### Imputation of sampling probability

The vector sampling *s* has two parts: the observed part *s^o^* and the unobserved part *s^u^*. Of the 1130 total observations, 539 have a duration of 3 or more and have an observed gap statistic. The gap statistic for the remaining 591 observations was imputed. As stated above, the unobserved part is then imputed, or filled in, based on the observed part *s^o^*. Because sampling varies between 0 and 1, I chose to model it as a Beta regression with matrix *W* being a *N ×* (*D −* 3) matrix of covariates (i.e. geographic range size, environmental preference, body size; no interactions). Predicting sampling probability using the other covariates that are then included in the model of duration is acceptable and appropriate in the case of imputation where the sample goal is accurate prediction (Gelman and Hill, 2007; Rubin, 1996). Not including these covariates can lead to biased estimates of the imputed variable; if the covariates themselves are related, not including them will bias this correlation towards zero which then leads to improper imputation and inference (Rubin, 1996).

The Beta regression is defined

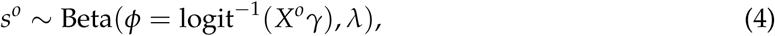

where γ is a length *D* vector of regression coefficients and *X* is defined as above. The Beta distribution used in the regression is reparameterized in terms of a mean parameter

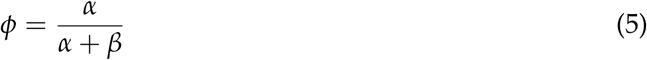

and total count parameter

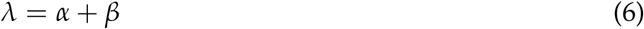

where α and β are the characteristic parameters of the Beta distribution (Gelman et al., 2013).

The next step is to estimate *s^u |^s^o^*, *X^o^*, *X^u^*, γ which is used as a covariate of taxon duration (Eq. 2). All the elements of γ, δ (Eq. 2), and λ (Eq. 4) were given weakly informative priors as recommended by Team (2017):

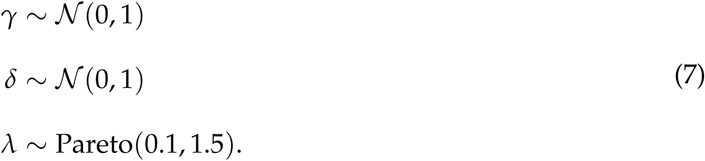

The imputed values are estimated simultaneously and in the same manner as all other parameters; this ensures that all uncertainty surrounding these unobservable covariate values is propagated through to all estimates.

### Posterior inference and posterior predictive checks

The joint posterior was approximated using a Markov-chain Monte Carlo routine that is a variant of Hamiltonian Monte Carlo called the No-U-Turn Sampler (Hoffman and Gelman, 2014) as implemented in the probabilistic programming language Stan (Stan Development Team, 2014). The posterior distribution was approximated from four parallel chains run for 40,000 steps, split half warm-up and half sampling and thinned to every 20th sample for a total of 4000 posterior samples. Starting conditions for sampling were left at the CmdStan defaults for interface except for the following changes: adapt delta was set 0.95 to ensure no divergent samples, and initial value was set to 0 which allows for stable initial samples. Posterior convergence was assessed using both standard MCMC and HMC specific diagnostics: scale reduction factor 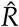, effective sample size or ESS, energy (target *>* 0.2), presence and number of divergent samples, and number of samples that saturated the maximum trajectory length. For further explanation of these diagnostic criteria, see the Stan Manual (Team, 2017).

After the model was fitted to the data, 100 datasets were simulated from the posterior predictive distribution of the model. These simulations were used to test for adequacy of model fit as described below.

Survival analysis is complicated by censored observations, where the ultimate time of extinction for some taxa could not be fully observed during the study window. Posterior predictive simulations for these observations must be similarly censored. To accomplish this, posterior predictive simulated durations for right-censored observations where the minimum of its final observed duration and the simulated duration. For left-censored individuals, their simulated duration was set to a minimum of one stage.

Model adequacy was evaluated using a series of posterior predictive checks. Posterior predictive checks are a means for understanding model fit or adequacy. Model adequacy means that if our model is an adequate descriptor of the data, then data simulated from the posterior predictive distribution should be similar to the observed given the same covariates, etc. (Gelman et al., 2013). Posterior predictive checks generally compare some property of the empirical data to that property estimated from each of the simulated datasets. Additionally, for structured datasets like the one analyzed here, the fit of the model to different parts of the data can be assessed, which in turn can reveal a great deal if the model has good fit to some aspects of data but not others; this is when when knowledge of the biological, geological, or paleoenvironmental context of the data becomes important in order to explain what unmodeled processes might lead to these discrepancies between our data and the model (Gelman et al., 2013).

The types of posterior predictive tests used in this analysis fall into two categories: comparison of observed mean and median genus duration to a distribution of mean and median genus duration estimates from the posterior simulations; and comparison of a non-parametric estimate of the survival function from the observed data to estimates of that same survival function from the simulations. These posterior predictive tests were done for the entire dataset as a whole and for each of the origination cohorts individually.

The survival function describes the probability of a taxon persisting given that it has survived up to time *t*; this is expressed *P*(*T ≥ t*) because *T* is the true extinction time of the species and *t* is some arbitrary time of observation and we are estimating that probability that *t* is less than *T*. It is important to note, however, that the survival function does not reflect density of observations unlike e.g. histograms. Instead, this posterior predictive check reflects the model’s ability to predict genus survival.

All code necessary to reproduce this analysis are available through an archived Zenodo repository DOI https://zenodo.org/record/1402252. Additionally, this project is hosted at https://github.com/psmit

## Results

I first present the results of the multiple posterior predictive checks for the whole dataset as well as each of the origination cohorts. I next present the parameter posterior estimates and their interpretations.

Comparisons between the observed distribution of durations to the distributions of 100 simulated datasets reveals the relatively good but heterogeneous fit of the model to the data (Fig. 1). The two major aspects of possible misfit that are observable are at durations of 2-3 stages. The model slightly under-estimates the number of observations with duration of 2 or 3 stages. The goal of this model is estimating the expected duration of a genus given its covariate information. While the model estimates are not exact, it is possible that our model fits the bulk of our data well but fits poorly towards the extreme values.

**Figure 1:**
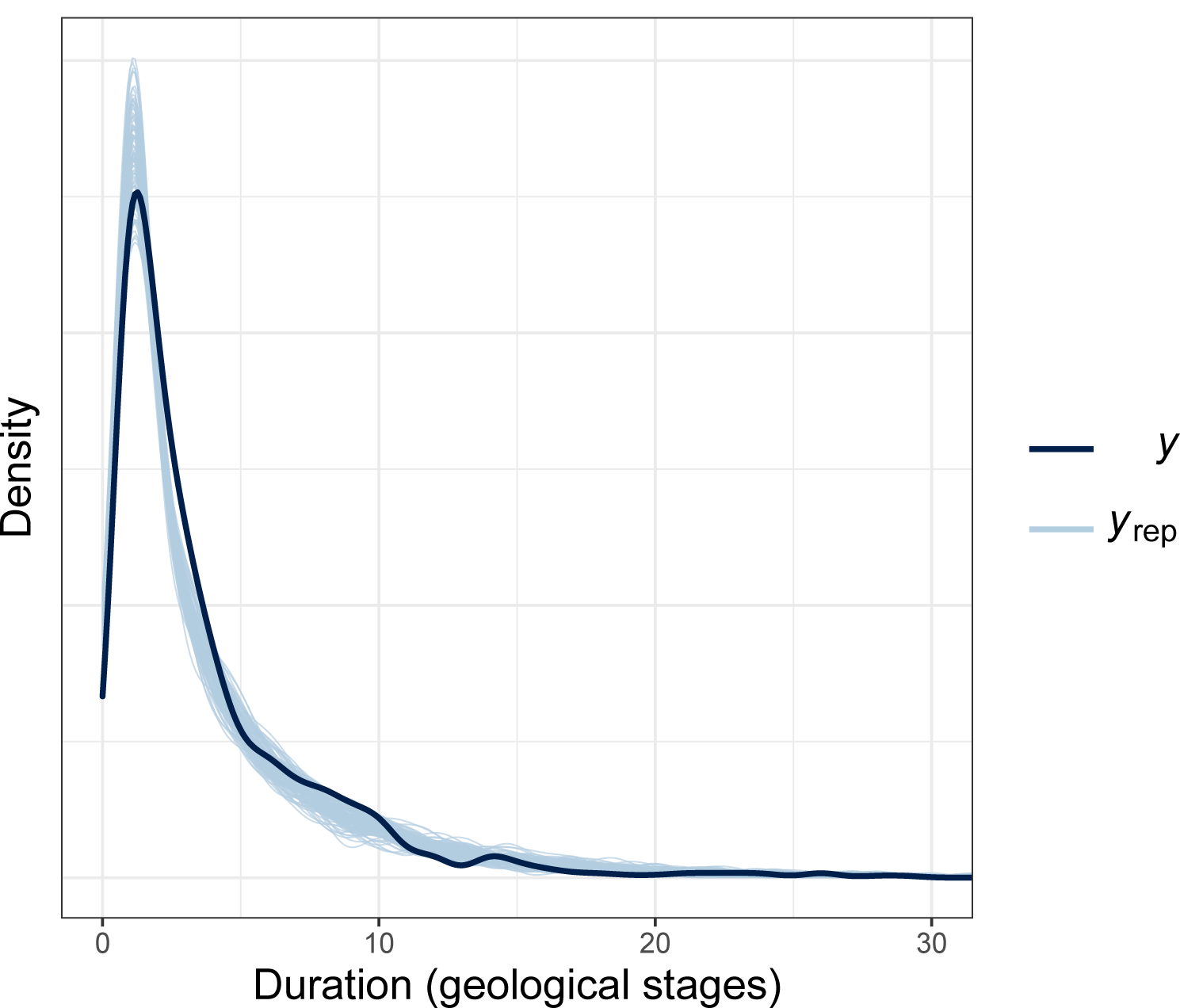
Comparison of the distribution of the observed data (black) to 100 simulated distributions (blue). This is a close-up view of the bulk of the distribution which shows the more subtle aspects of (mis)fit between the data and the model.

The similarity of the empirical data and from 100 simulated datasets provides a more nuanced picture of model adequacy (Fig. 2). The survival curves of the 100 simulated datasets are very similar to the survival function estimated from the empirical data. The major points of misfit between the model and the data are taxa with duration 1 stage, and taxa with a duration at least 10-13 stages. The major divergence between the observed and the estimated applies to taxa with a less than 15% probability of continuing to survive.

**Figure 2:**
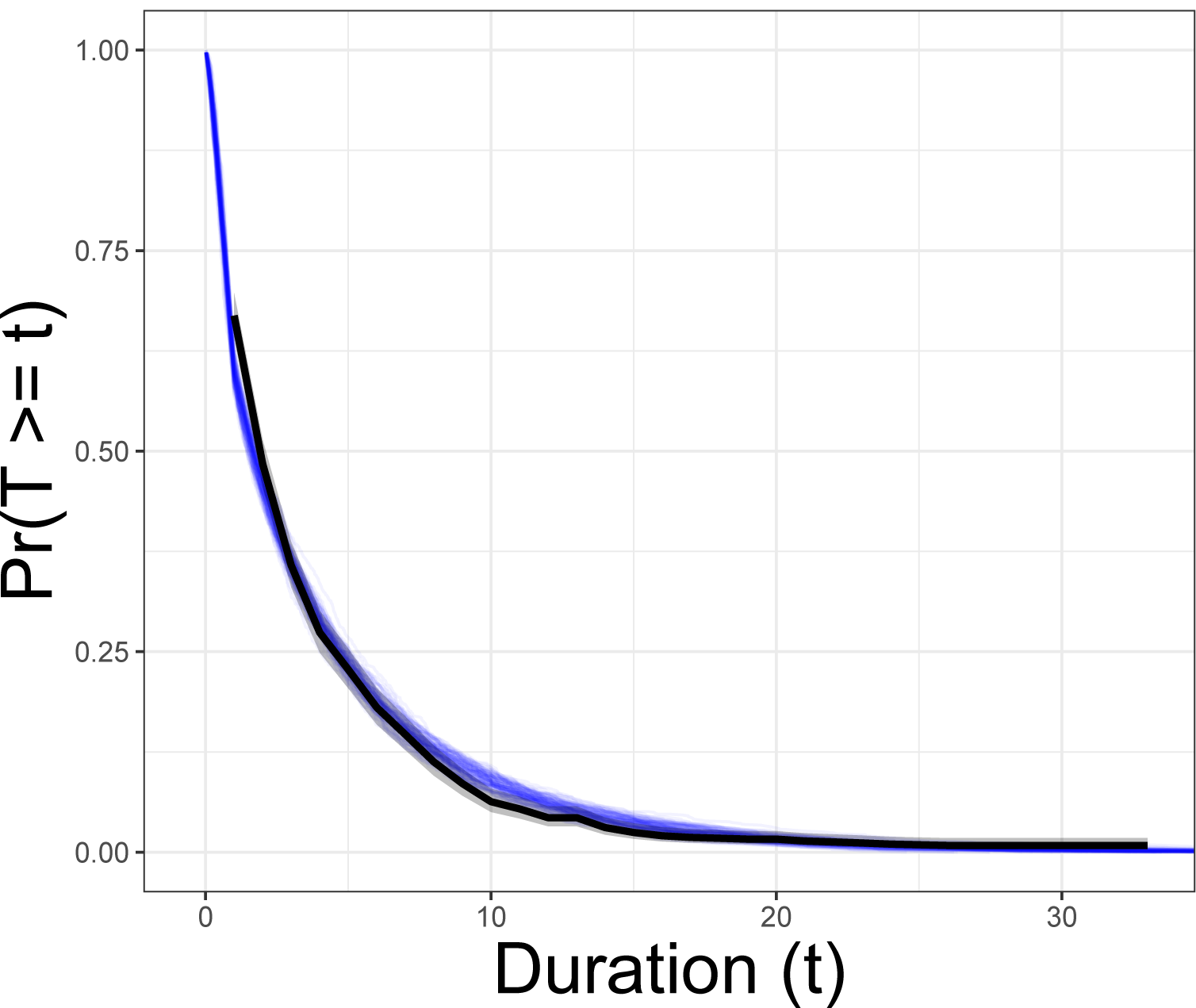
Comparison of the empirical estimate of *S*(*t*) (blue) versus estimates from 100 posterior predictive data sets (black). *S*(*t*) corresponds to the probability that the age of a genus *t* is less than the genus’ ultimate duration *T*.

Model adequacy at the total data level was assessed through comparison of the mean and median of the observed data to those from simulated data sets. While the previous posterior predictive checks have focused on the relatively good but heterogeneous fit of the model to the entire distribution of the data, the fitted model’s ability to predict the mean and median of the observed data appears adequate (Fig. 3, 4). Because the principle goal of this model is to obtain adequate prediction of a taxon’s expected duration for a given set of ecological covariates, the seemingly adequate fit of our model to mean taxon duration is reassuring (Fig. 3). Additionally, given the skewness of the observed taxon durations (Fig. 1), the ability for the model to recapitulate the median observed taxon duration points to the overall good fit of the model to the data.

**Figure 3:**
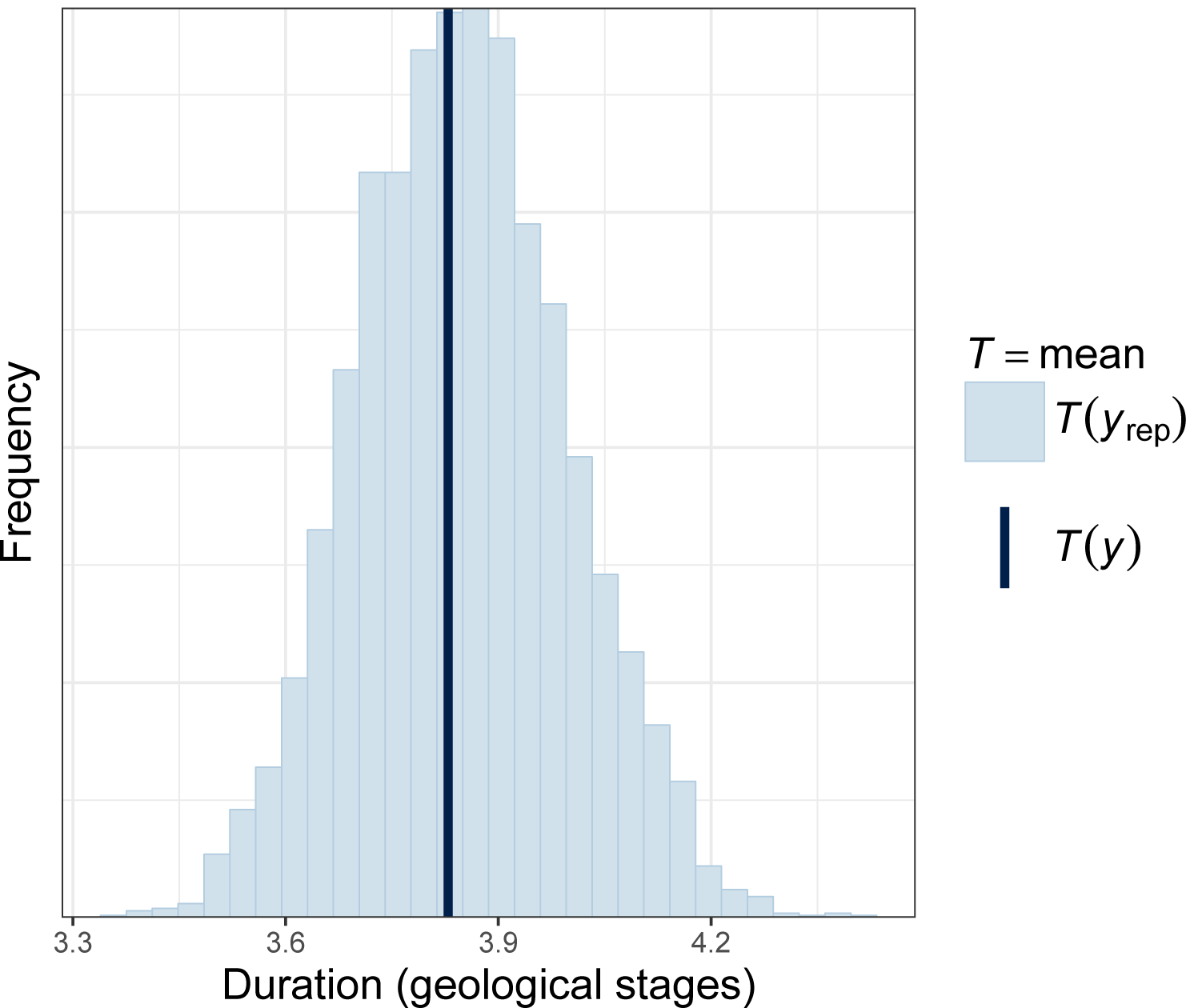
Comparison of the (A) observed mean genus duration (black vertical line) to a distribution of means estimated from 100 simulated datasets (blue). Model fit is evaluated by the similarity between the observed and the estimated, where good fit is demonstrated by the vertical line being “within” the simulated distribution.

**Figure 4:**
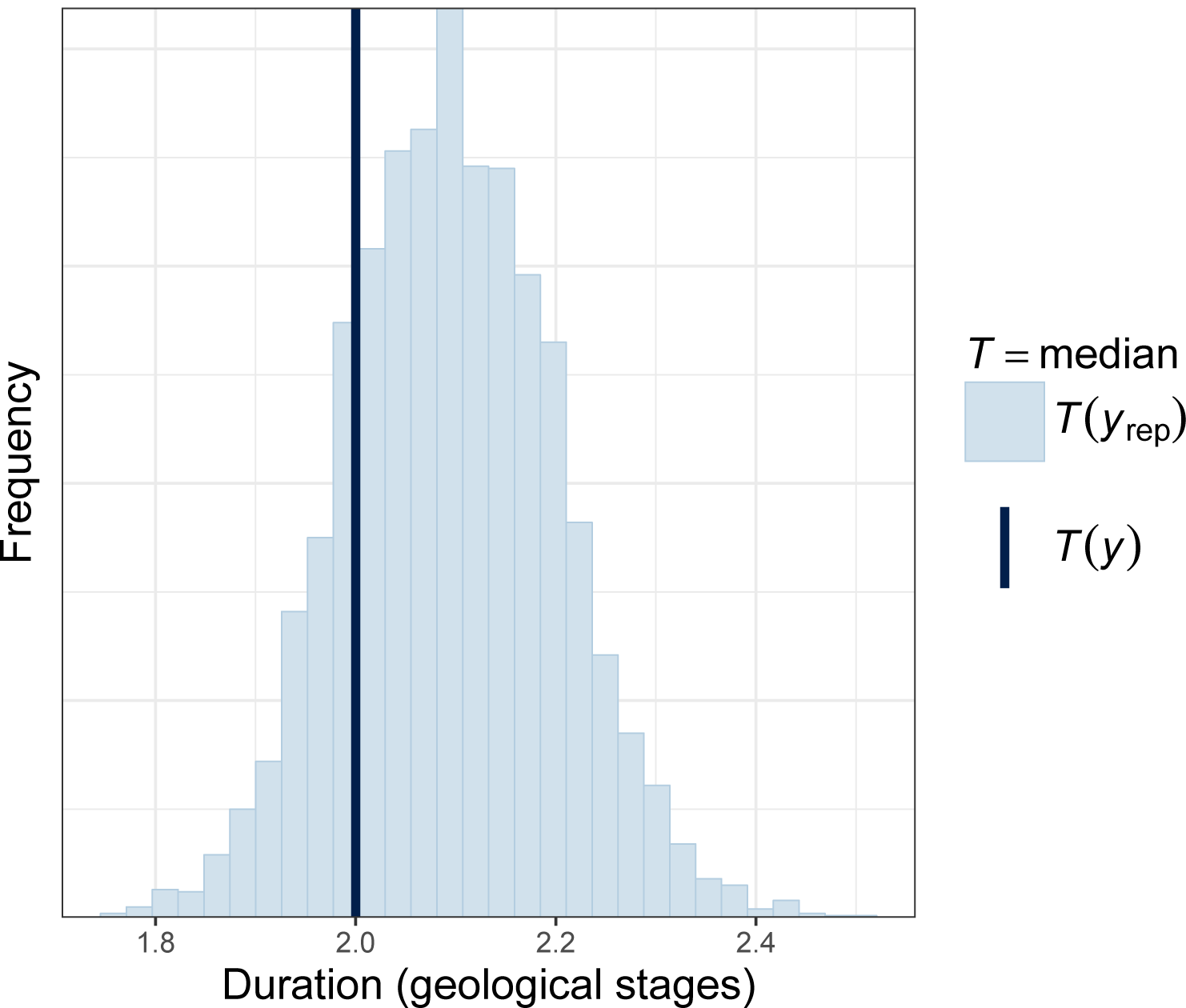
Comparison of the observed median genus duration (black vertical line) to a distribution of medians estimated from 100 simulated datasets (blue). Model fit is evaluated by the similarity between the observed and the estimated, where good fit is demonstrated by the vertical line being “within” the simulated distribution.

When considered together, all of the above posterior predictive checks indicate adequate model fit for key questions such as expected taxon duration (Fig. 3). However, there is obviously heterogeneity in model fit because, while the model can recapitulate some aspects of the observed data (Fig. 3, 4), there are obvious discrepancies between the model and the data (Fig. 1, 2). By performing the same posterior predictive tests for each of the origination cohorts, it may be possible to get a better picture of the sources of model misfit.

When the posterior predictive tests are visualized for each of the origination cohorts, a complex picture of model fit emerges. For nearly every origination cohort, the model is able to recapitulate the observed mean duration (Fig. 5). In comparison, the model has a much more heterogeneous fit to each origination cohort’s median taxon duration (Fig. 6). The skewness of the distribution underlying figure 1 means that for some origination cohorts, median duration might be pegged at 1 stage; this means that the posterior predictive distributions for some cohorts can be extremely skewed.

**Figure 5:**
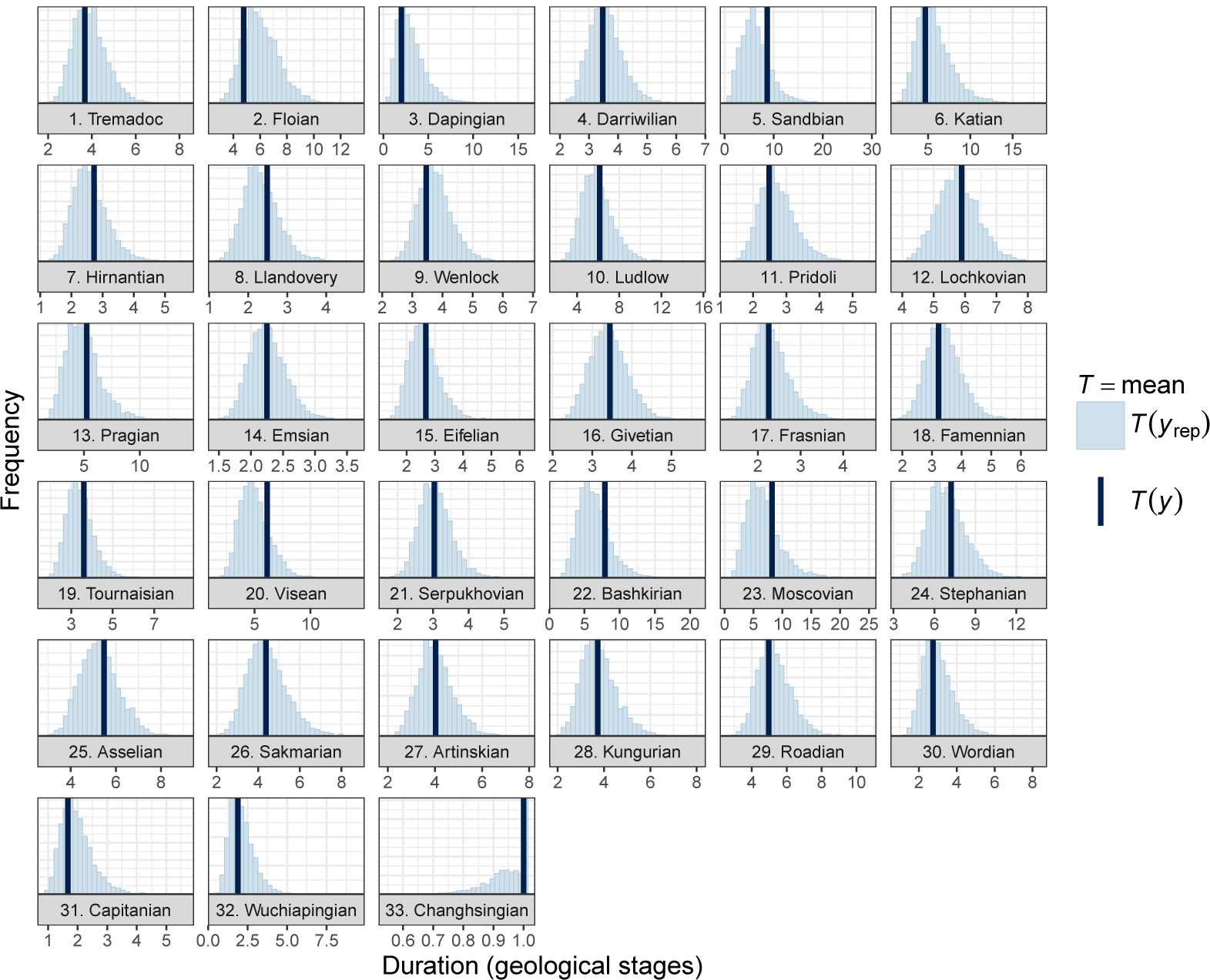
Comparison of the observed mean genus duration (black vertical line) to a distribution of means estimated from 100 simulated datasets (blue) for each of the origination cohorts. Model fit is evaluated by the similarity between the observed and the estimated, where good fit is demonstrated by the vertical line being “within” the simulated distribution.

**Figure 6:**
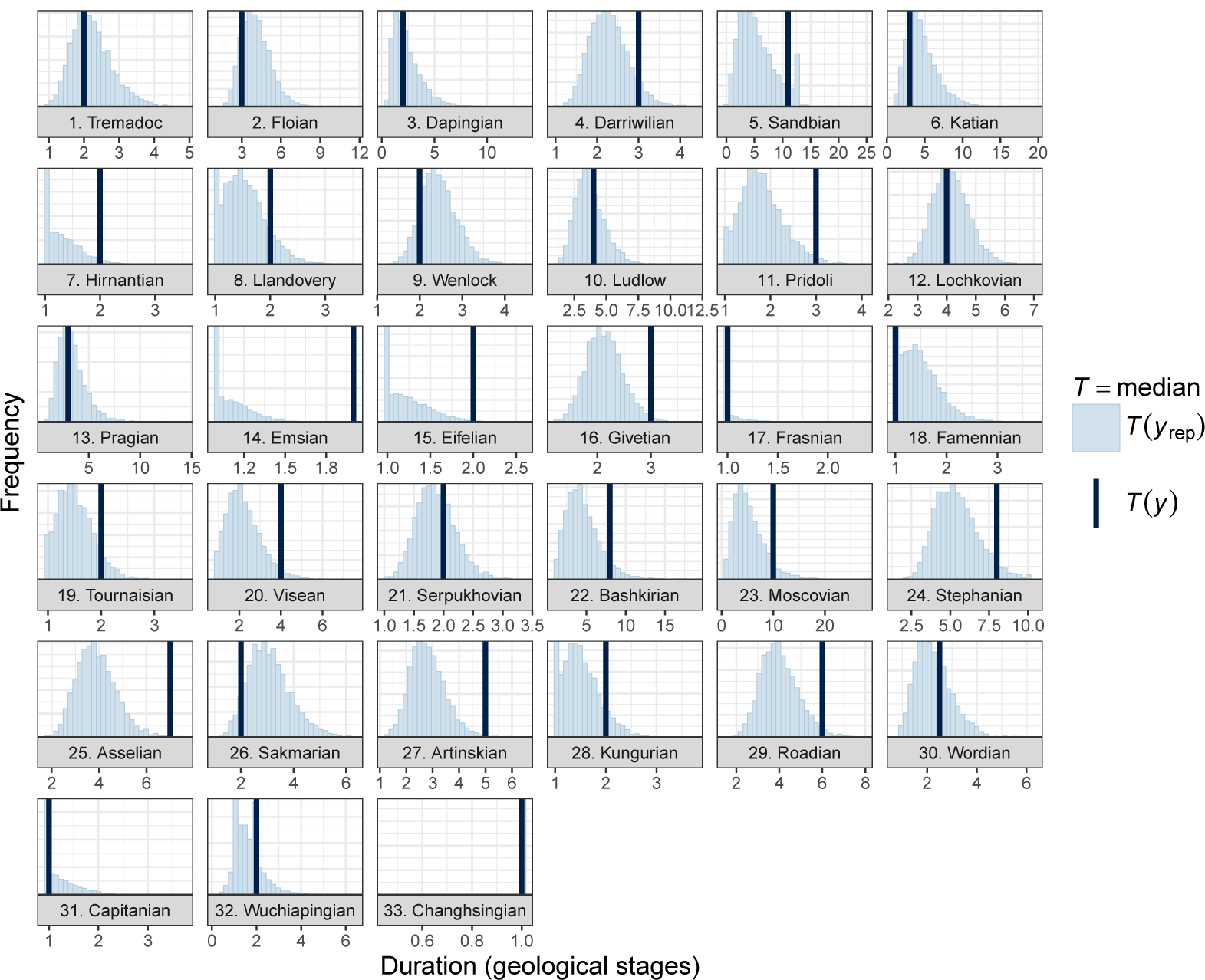
Comparison of the observed median genus duration (black vertical line) to a distribution of medians estimated from 100 simulated datasets (blue) for each of the origination cohorts. Model fit is evaluated by the similarity between the observed and the estimated, where good fit is demonstrated by the vertical line being “within” the simulated distribution.

These results indicate that this model is very good at recapitulate mean taxon duration (Fig. 3, 5) and that it is capable of estimating overall median duration and median duration of most origination cohorts (Fig. 4, 6). The poor model fit to some origination cohorts may indicate that these cohorts are undergoing a different extinction process whose aspects are unmodeled in this analysis. For those cohorts where the model recapitulates the empirical survival function, the model is capturing some aspect of the processes underlying taxon extinction.

A larger than average geographic range is expected to have a positive effect on taxon survival (Table 1). The cohort-level estimate of the effect of geographic range size indicates that as a taxon’s geographic range increases, that taxon’s duration is expected to increase (Table 1). Given the estimates of µ^*r*^ and τ^*r*^, there is an approximately 3.7% (±4.3% SD) probability that this relationship would be reversed (Pr*N* (µ^*r*^, τ^*r*^) *>* 0)).

**Table 1:** Estimates of group-level and invariant parameter values for the fitted model analyzed here.

| Category | Parameter | Effect of... | Mean | SD | 10% | 50% | 90% |
| --- | --- | --- | --- | --- | --- | --- | --- |
| Mean | $\mu^i$ | intercept | -3.04 | 0.19 | -3.29 | -3.04 | -2.80 |
| | $\mu^r$ | geographic range | -0.98 | 0.16 | -1.17 | -0.98 | -0.78 |
| | $\mu^v$ | environmental preference | -0.76 | 0.18 | -0.99 | -0.76 | -0.53 |
| | $\mu^{v^2}$ | environmental preference<sup>2</sup> | 3.15 | 0.35 | 2.71 | 3.15 | 3.59 |
| | $\mu^m$ | body size | -0.02 | 0.12 | -0.17 | -0.02 | 0.14 |
| Standard deviation | $\tau^i$ | intercept | 0.50 | 0.11 | 0.38 | 0.50 | 0.65 |
| | $\tau^r$ | geographic range | 0.49 | 0.16 | 0.29 | 0.49 | 0.70 |
| | $\tau^v$ | environmental preference | 0.83 | 0.16 | 0.63 | 0.82 | 1.05 |
| | $\tau^{v^2}$ | environmental preference<sup>2</sup> | 1.49 | 0.35 | 1.08 | 1.46 | 1.94 |
| | $\tau^m$ | body size | 0.47 | 0.12 | 0.32 | 0.46 | 0.63 |
| Other | $\delta$ | sampling | 0.90 | 0.15 | 0.71 | 0.89 | 1.08 |
| | $\alpha$ | ageing | 1.36 | 0.04 | 1.30 | 1.36 | 1.42 |

Note: These parameters are the group-level estimates of the effects of biological traits on brachiopod generic survival, the standard deviation of the between-cohort effects, as well as the estimates of the effect of sampling $\delta$ and the Weibull shape parameter $\alpha$ . The mean, standard deviation (SD), 10th, 50th, and 90th quantiles of the marginal posteriors are presented.

Body size measured as valve length is estimated to have no effect on duration for most of the post-Cambrian Paleozoic (Table 1).

The group-level relationship between effect of environmental preference and duration is estimated from µ^*v*^ and 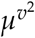. The estimate of µ^*v*^ indicates that taxa which slightly prefer epicontinental environments to open-ocean environments are expected to have a greater duration than open-ocean favoring taxa (Table 1). Additionally, given the estimate of between-cohort variance τ^*v*^, there is approximately 18.1% (±7.5% SD) probability that, for any given cohort, taxa which favor open-ocean environments would have a greater expected duration than taxa which favor epicontinental environments (Pr(*N* (µ^*v*^, τ^*v*^) *>* 0)). The estimate of 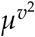 indicates that the overall relationship between environmental preference and log(σ) is concave-down (Fig. 7), with only a 2.5% (±2.9% SD) probability that any given cohort is convex up 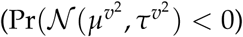.

**Figure 7:**
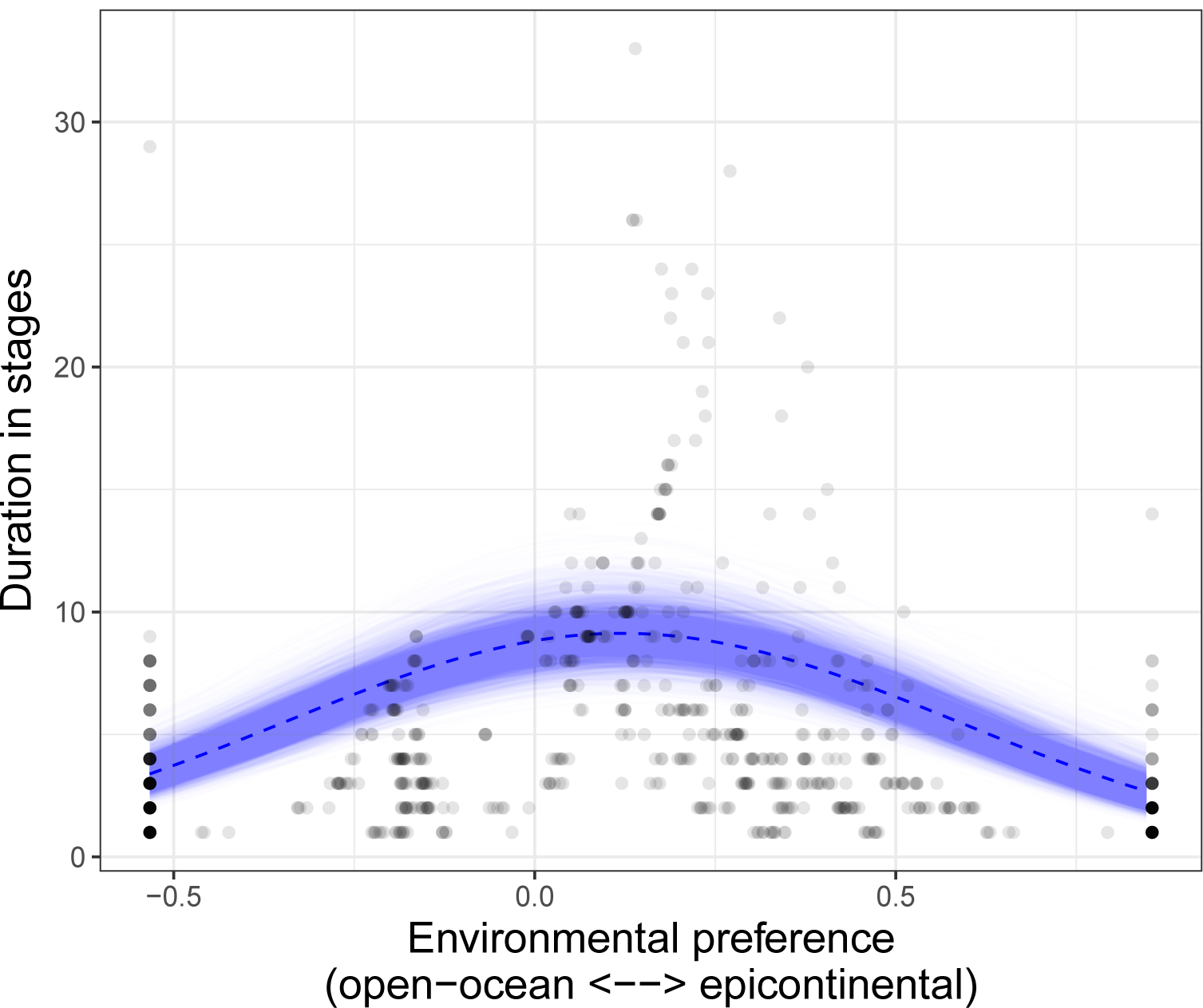
The overall expected relationship between environmental affinity *v_i_* and a log(σ) when r = 0 and m = 0. The 1000 semi-transparent lines corresponds to a single draw from the posterior predictive distribution, while the highlighted line corresponds to the median of the posterior predictive distribution. The overall relationship demonstrates a greater durations among environmental generalists than specialists. Additionally, because the apex of is rightward from 0, taxa favoring epicontinental environments are expected to have a slightly longer durations than those favoring open-ocean environments. The tick marks along the bottom of the plot correspond to the (rescaled) observed values of environmental preference.

The cohort-specific relationships between environmental preference and log(σ) were calculated from the estimates of β^0^, β^*v*^, and 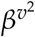 (Fig. 8) and reflect how these three parameters act in concert, not individually (Fig. 9). Because of the relationship between β^*v*^ and 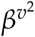, it is important to consider them together when drawing conclusions from the model. In many cases, the cohort-specific estimated relationship between environmental preference and duration is approximately equal to the group-level average, but for 14 of the 33 analyzed origination cohorts at least one of these three parameters are noticeably different from the group-level average.

**Figure 8:**
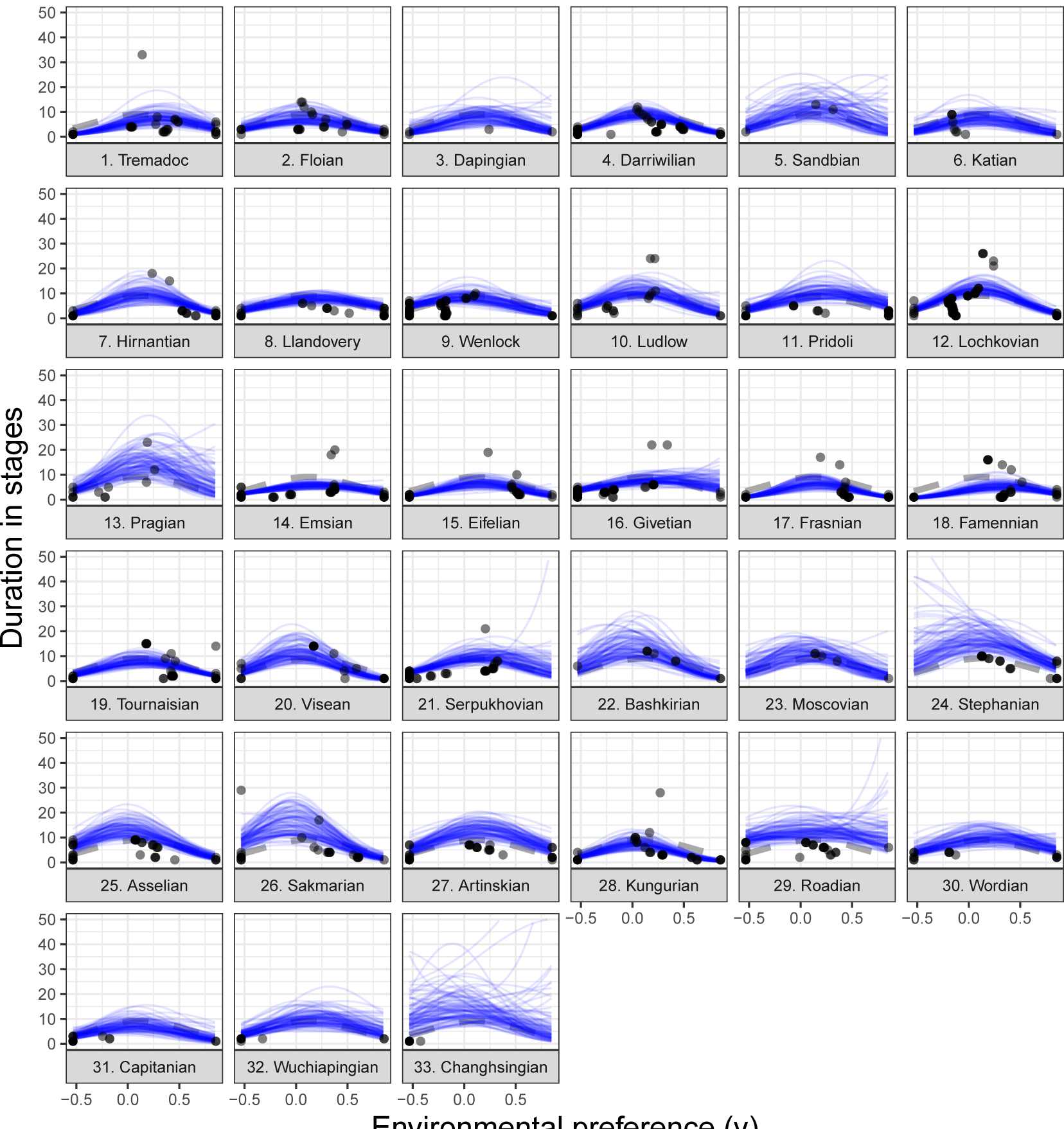
Comparison of origination cohort-specific (posterior predictive) estimates of the effect of environmental preference on log(σ) to the mean overall estimate of the effect of environmental preference. Cohort-specific estimates are from 100 posterior predictive simulations across the range of (transformed and rescaled) observed values of environmental preference. The oldest cohort is in the top-left and younger cohorts proceed left to right, with the youngest cohort being the right-most facet of the last row. Panel names correspond to the name of the stage in which that cohort originated.

**Figure 9:**
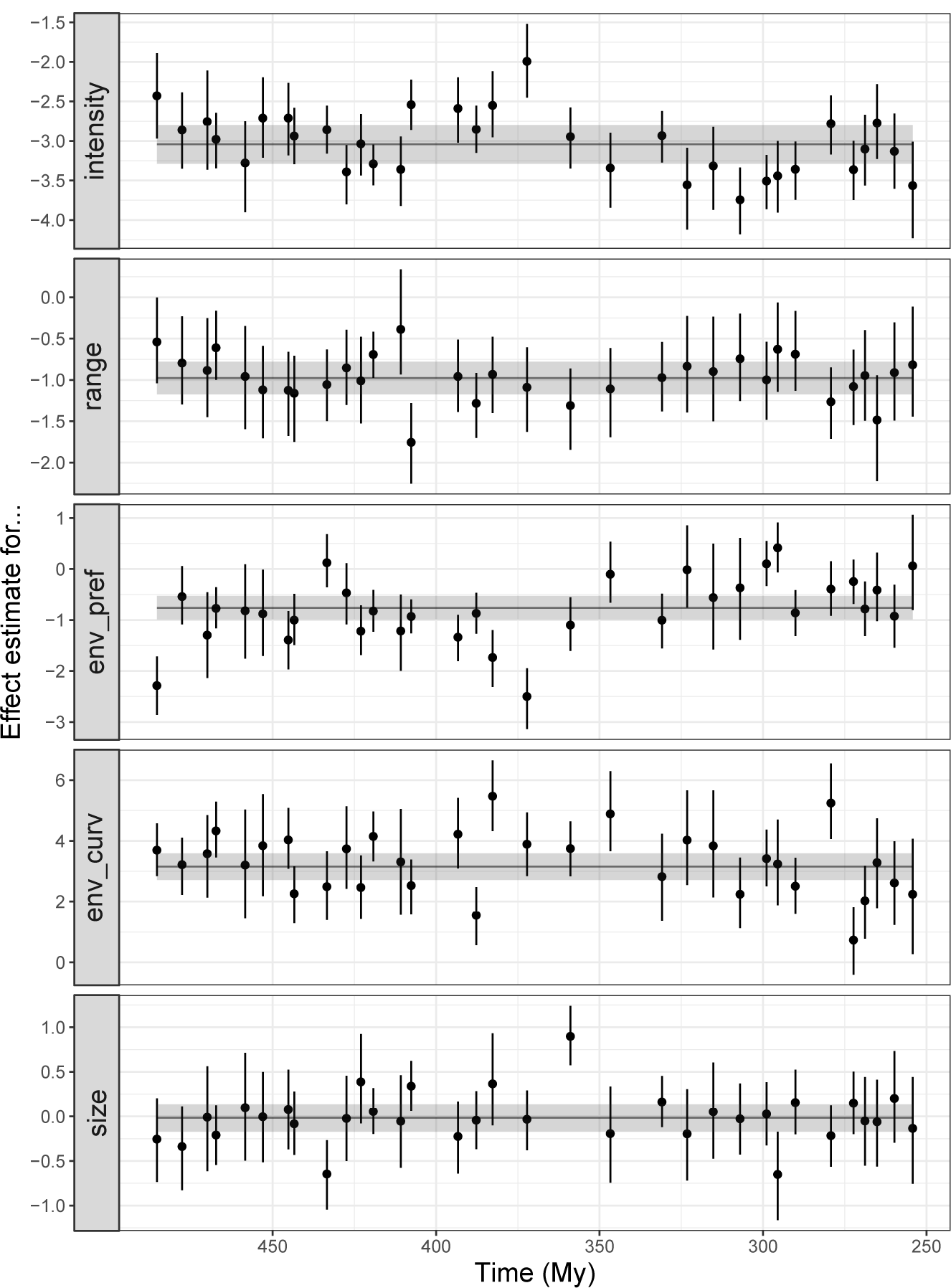
Comparison of cohort-specific estimates of β^0^, the effect of geographic range on extinction risk β^*r*^, the effect of environmental preference β^*v*^ and 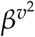, and body size β^*m*^. Points correspond to the median of the cohort-specific estimate, along with 80% credible intervals. Points are plotted at the midpoint of the cohorts stage of origination in millions of years before present (My). Black, horizontal lines are the overall estimates of covariate effects along with 80% credible intervals (shaded).

There is an approximately 90.4% probability that cohort estimates of β^0^ and β^*r*^ are negatively correlated, with median estimate of correlation being −0.35 (Fig. 10). This result means that for any cohort, we would expect that if extinction intensity increases (β^0^ increases), the effect of geographic range on duration increases (β^*r*^ decreases). This result is strong evidence for a relationship between intensity and selectivity with respect to geographic range size.

**Figure 10:**
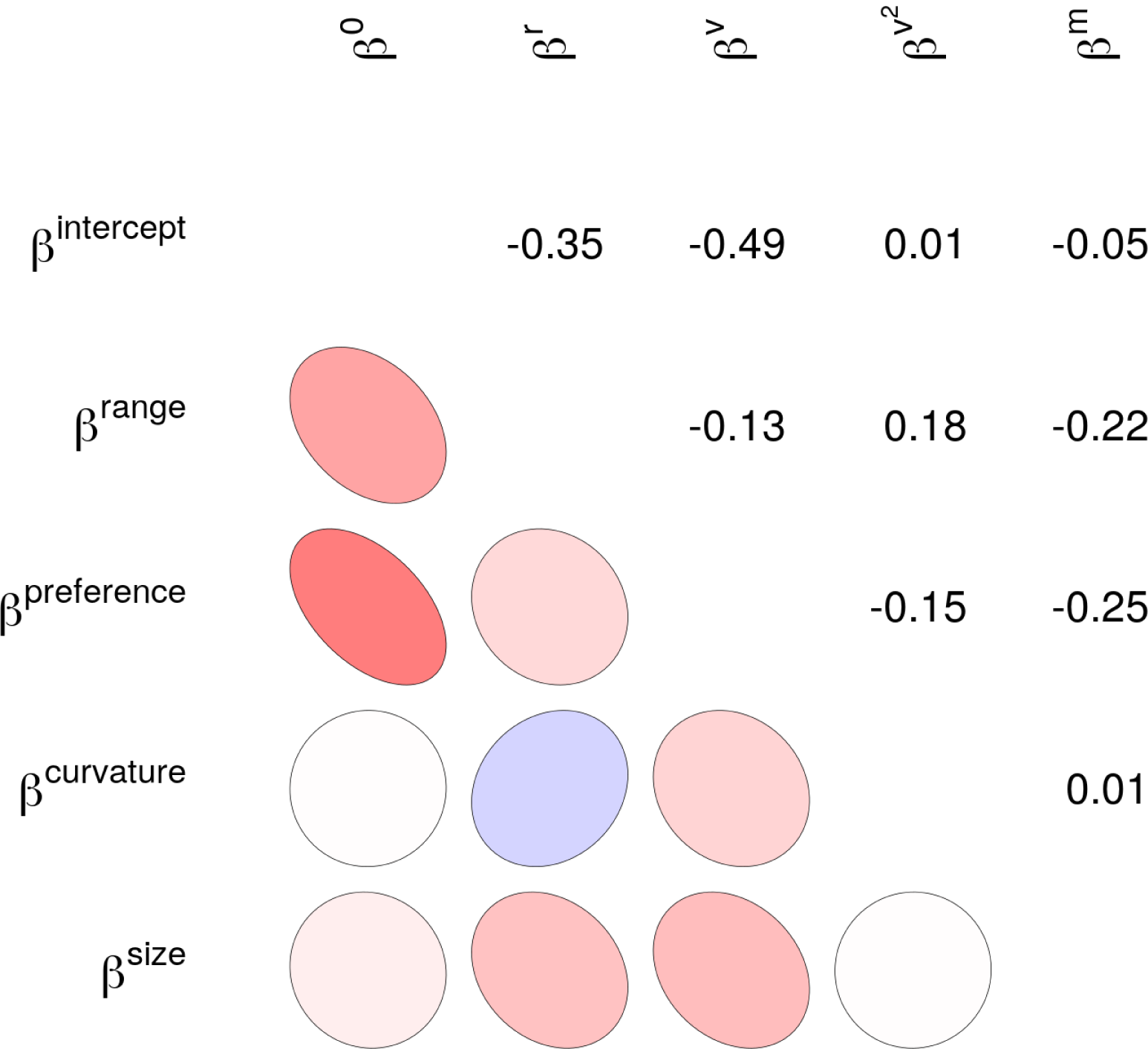
Mixed graphical and numerical representation of the correlation matrix Ω of variation in cohort-specific covariate estimates. These correlations are between the estimates of the cohort-level effects of covariates, along with intercept/baseline extinction risk. The median estimates of the correlations are presented numerically (upper-triangle) and as idealized ellipses representing that much correlation (lower-triangle). The darkness of the ellipse corresponds to the magnitude of the correlation.

I estimate a 97.9% probability that the cohort-specific estimates of β^0^ and β^*v*^ are negatively correlated, with a median correlation of −0.49 (Fig. 10). This result means that as extinction intensity increases it is expected that epicontinental taxa become more favored over open-ocean environments (i.e. as β^0^ increases, β^*v*^ decreases). This result is strong evidence for a relationship between intensity and selectivity with respect to the linear aspect of environmental preference.

Correlations between the non-intercept estimates reflect potential similarities in selective pressures between cohorts, however there is only weak evidence of any potential cross-correlations in cohort-specific covariate effects(Fig. 10). There is an approximate 31.2% probability that β^*r*^ and β^*v*^ are positively correlated. This lack of cross-correlation may be due in part to the higher between-cohort variance of the effect of environmental preference τ^*v*^ compared to the very small between-cohort variance in the effect of geographic range τ^*r*^ (Table 1); the effect of geographic range might simply not vary enough relative to environmental preference.

Conversely, there is a 74.6% probability that estimates of the effect of geographic range (β^*r*^) and the quadratic aspect of environmental preference 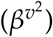 are positively correlated; this is weak evidence of a relationship between the effects of these covariates (Fig. 10). Thus, as the effect of geographic range increases, we might expect the peakedness of relationship between environmental preference and duration to increase. However, because there is only a 74.6% probability of a positive correlation, this result cannot interpreted with authority. Instead, this result is an opportunity for future research to further explore the potential relationship between geographic range, environmental preference, and species duration.

Sampling was found to have a negative effect (positive δ) on duration: greater sampling, shorter duration (Table 1). While potentially counter-intuitive, this result is most likely due to some long lived taxa only be sampled in the stages of it’s first and last appearance. Also, longer lived taxa have more opportunities to evade sampling than shorter lived taxa. These two factors will lead to this result.

The Weibull shape parameter α was found to be approximately 1.41 (±0.05 SD) with a 100% probability of being greater than 1. This result is not consistent with the Law of Constant Extinction (Van Valen, 1973) and is instead consistent with accelerating extinction risk with taxon age. This result is consistent with recent empirical results and may be caused by newly originating species having a fundamentally lower risk of extinction compared to species which have already originated (Quental and Marshall, 2013; Smits, 2015; Wagner and Estabrook, 2014). This result is also consistent with a recently proposed nearly-neutral evolution where competition/selection/evolution drives whole communities to increase in average fitness over time while still maintaining constant relative fitness across the community, thus older species are more likely to go extinct because of having a fundamentally lower average fitness than newly originating species (Rosindell et al., 2015). This result, however, is not consistent with other empirical results from the marine fossil record (Crampton et al., 2016; Finnegan et al., 2008) and could potentially be caused by the minimum resolution of the fossil record (Sepkoski, 1975). It is thus unclear whether a strong biological inference can be made from this result, which means that further work is necessary on the effect of taxon age on extinction risk.

## Discussion

The generating observation behind this study was that for bivalves at the end Cretaceous mass extinction event, the only biological trait that was found to affect extinction risk was geographic range, while traits that had previously been associated with difference in duration had no effect (Jablonski, 1986). This observation raises two linked questions: how does the effect of geographic range change with changing extinction intensity, and how to the effects of other biological traits change with changing extinction intensity?

I find that as intensity increases (β^0^ increases), the magnitude of the effect of geographic range increases (β^*r*^ decreases). I also find that as intensity increases, the difference in survival for taxa favoring epicontinental environments over open-ocean environments is expected to decrease; this is consistent with the results of Miller and Foote (2009). Finally, there is no evidence for a correlation between the effects of geographic range and environmental preference on taxon duration.

I find consistent support for the “survival of the unspecialized,” with respect to epicontinental versus open-ocean environmental preference, as a time-invariant generalization of brachiopod survival (Simpson, 1944). Taxa with intermediate environmental preferences are expected to have lower extinction risk than taxa specializing in either epicontinental or open-ocean environments (Fig. 7), though the curvature of the relationship varies from rather shallow to very peaked (Fig. 8). However, this relationship is not symmetric about 0, as taxa favoring epicontinental environments are expected with approximately 75% probability to have a greater duration than taxa favoring open-ocean environments. This description of environment preference is only one major aspect of a taxon’s environmental context, with factors such as bathymetry and temperature being further descriptors of a taxon’s adaptive zone (Harnik, 2011; Harnik et al., 2012; Heim and Peters, 2011; Nürnberg and Aberhan, 2013); inclusion of these factors in future analyses would potentially improve our understanding of the extent and complexity of the “survival of the unspecialized” hypothesis as it applies to all dimensions of an adaptive zone.

Hopkins et al. (2014), in their analysis of niche conservatism and substrate preference in marine invertebrates, found that brachiopods were among the least “conservative” groups, with taxa found to change substrate preference on short time scales. While substrate preference is not the same as environmental preference (as defined here), a question does arise: are there three classes of environmental preference instead of two? These classes would be taxa with broad tolerance (“true” generalists), inflexible specialists (“true” specialists), and flexible specialists. A flexible taxon is one with a narrow habitat preference at one time but with preference that changes over time with changing environmental availability. My analysis assumes that traits are constant over the duration of the taxon, meaning that this scenario is not detectable: taxa with broad tolerances and flexible taxa with narrow per-stage preference end up being treated the same way. Future work should explore how environmental preference changes over lineage duration in relation to environmental availability to estimate if the generalists–specialists continuum is actually a ternary relationship.

The analysis presented in this paper is an example of how to approach the interplay between selection and intensity using a continuous-survival framework. An alternative framework would be a discrete-time survival analysis (Tutz and Schmid, 2016) where species survival is tracked at discrete intervals. An example of a discrete-time survival approach that has become increasingly popular in paleontological analysis is the Cormack-Jolly-Seber (CJS) model (Liow et al., 2008; Liow and Nichols, 2010; Royle and Dorazio, 2008; Tomiya, 2013). Discrete-survival analysis has some advantages over continuous-time approaches, specifically the ease of including time-varying covariates and well known extensions for allowing incomplete sampling (e.g. CJS model).

Something that has not been modeled in these discrete-time analysis is the effect of an age-based varying-intercept or covariate on duration as recommded by Tutz and Schmid (2016); this is extremely important for estimating the effect of taxon age on survival. Those varying-intercept estimates would then be equivalent to the hazard function when all covariates are equal to 0 (Tutz and Schmid, 2016). A good avenue for future applied research would be a CJS-type model with survival modeled as a multi-level regression as in this study, combined with an age-based varying-intercept as recommended by Tutz and Schmid (2016). A major hurdle to this analysis would be the necessity of imputing all time-varying covariates for every taxon that is estimated to be present in a time intervals but was not sampled.

The model used here could be improved through either increasing the number of analyzed traits, expanding the hierarchical structure of the model to include other major taxonomic groups of interest, or including explicit phylogenetic relationships between the taxa in the model as an additional hierarchical effect. An example trait that may be of particular interest is the affixing strategy or method of interaction with the substrate of the taxon, which has been found to be related to brachiopod survival where, for cosmopolitan taxa, taxa that are attached to the substrate are expected to have a greater duration than those that are not (Alexander, 1977).

It is theoretically possible to expand this model to allow for comparisons within and between major taxonomic groups, which would better constrain the brachiopod estimates while also allowing for estimation of similarities and differences in cross-taxonomic patterns. The difficulty with this particular model expansion is in finding a similarly well-sampled taxonomic group that is present during the Paleozoic. Potential groups include Crinoidea, Ostracoda, and other members of the “Paleozoic fauna” (Sepkoski, 1981).

With significant updates, it would also be possible to compare the brachiopod record with Moden groups such as bivalves or gastropods (Sepkoski, 1981), while remembering that the groups may not necessarily share all cohorts with the brachiopods. This particular model expansion would act as a test of any universal cross-taxonomic patterns in the effects of emergent traits on extinction, as has been proposed for geographic range (Payne and Finnegan, 2007). Additionally, this expanded model would also act as a test of the distinctness of the Sepkoski (1981) three-fauna hypothesis in terms of trait-dependent extinction.

Traits like environmental preference or geographic range (Hunt et al., 2005; Jablonski, 1987) are most likely heritable. Without phylogenetic context, this analysis assumes that differences in extinction risk between taxa are independent of the shared evolutionary history of those taxa (Felsenstein, 1985). In contrast, the origination cohorts only capture shared temporal context. For example, if taxon duration is phylogenetically heritable, then closely related taxa may have more similar durations as well as more similar biological traits. Without taking into account phylogenetic similarity the effects of these biological traits would be inflated solely due to inheritance. The inclusion of phylogenetic context as an additional individual-level hierarchical effect, independent of origination cohort, would allow for determining how much of the observed variability is due to shared evolutionary history versus shared temporal context versus actual differences associated with biological traits (Smits, 2015).

The combination and integration of the phylogenetic comparative and paleontological approaches requires both sources of data, something which is not possible for this analysis because there is no phylogenetic hypothesis for all Paleozoic taxa, which is frequently the case for marine invertebrates with a good fossil record. When both data sources are available the analysis can more fully address the questions of interest in macroevolution (Fritz et al., 2013; Harnik et al., 2014; Raia et al., 2012, 2013; Simpson et al., 2011; Slater, 2013, 2015; Slater et al., 2012; Smits, 2015; Tomiya, 2013).

## Conclusion

My analysis demonstrates that for post-Cambrain Paleozoic brachiopds, as extinction intensity increases and average fitness decreases, such as in a mass extinction, the trait-associated differences in fitness (selection) would increase and be greater than aeverage. In contrast, during periods of low extinction intensity when fitness is greater than average, my model predicts that geographic range – and environmental preference – associated with differences in fitness (i.e. selection) would decrease and be less than average. Taken together, these results point to a potential macroevolutionary mechanism behind differences in trait-based survival during mass extinctions due to a correlation between intensity and selectivity. Additionally, I find continued support for greater survival in environmental generalists over specialists; this is further evidence that the long standing “survival of the unspecialized” hypothesis (Baumiller, 1993; Liow, 2004, 2007; Nürnberg and Aberhan, 2013, 2015; Simpson, 1944, 1953; Smits, 2015) should be considered the default hypothesis. Overall, this analysis further refines our knowledge of brachiopod extinction dynamics while also revealing a potential macroevolutionary mechanism behind the difference between so-called mass and background extinction regimes.

